# A Singapore-centric Fungal Dataset of 518 Cultivated Strains with Visual Phenotypes and Taxonomic Identity

**DOI:** 10.1101/2025.06.15.659681

**Authors:** Darren Ten Wei Xian, Fong Tian Wong, Yee Hwee Lim, Winston Koh

## Abstract

The fungal kingdom represents a greatly untapped resource to produce a wide range of bioactive secondary metabolites, including antibiotics, anticancer agents, industrially significant dyes and enzymes. To-date, it is estimated only less than 5% of all fungi have been characterised, a deficit that is especially pronounced in tropical regions like Singapore, where fungal diversity remains underexplored compared to northern hemisphere counterparts. This underlines the urgency and importance of our research which motivated the creation of our curated dataset, aiming to address this gap and contribute to understanding the broader ecosystem. We developed a generalisable cultivation workflow that enables systematic strain preparation, supports high-resolution imaging, and yields sufficient fungal biomass amenable for genomic analyses. This resulted in a diverse collection of 518 phylogenetically and ecologically varied fungal strains from both terrestrial and marine environments in biodiverse Singapore. The curated dataset from this project captures both taxonomic identity and colony-level morphological traits serving as a foundation for visual phenotype to taxonomy mapping through the integration of computer vision.

## Background & Summary

Fungi comprise one of the most diverse and ecological indispensable kingdoms on Earth. They are deeply integrated into the evolutionary history of life and continually serve as primary decomposers, facilitating nutrient cycling that sustains ecosystems. (Case et al., 2022; Peay et al., 2016; Tedersoo et al., 2014). As one of nature’s most prolific chemists, fungi synthesise an extraordinary repertoire of molecules, with transformative applications across medicine, biotechnology and industry. (Bhattarai et al., 2021; Danner et al., 2023; Hyde et al., 2024; Pye et al., 2017). The vast potential and opportunities offered by fungal chemistry has pushed efforts to broaden the exploration of fungal diversity, as exemplified by several large-scale initiatives. Notably, the Joint Genome Institute (JGI)’s MycoCosm’s platform currently hosts over 2,500 publicly available fungal genomes, compiled through a combination of community-contributed datasets and in-house sequencing efforts. This is primarily from the 1000 Fungal Genomes Project, which prioritises sequencing of reference genomes across more than 500 fungal families (Ahrendt et al., 2023; Grigoriev et al., 2014). Another major initiative is MycoBank, a global database and inter-repository hub for the registration and standardisation of novel fungal species nomenclature, also providing reference access to associated fungal taxonomic and genomic resources (Crous et al., 2004; Robert et al., 2013).

Despite this global momentum to elucidate the projected existence of 5 to 12 million fungal species, only ∼150,000 species have been formally described, with an even smaller proportion explored for their biotechnological potential (Huberman, 2021; Hyde, 2022; Paterson et al., 2023; Wu et al., 2019). This disparity is particularly pronounced in tropical regions, where characterized fungi species are predominantly from the northern hemisphere (Aime & Brearley, 2012; Stallman et al., 2024). Situated in the biodiverse equatorial region of Southeast Asia, Singapore harbours a remarkable diversity of flora and fauna, with an estimated 40,000 species, with a recent biodiversity survey formally documented over 3,000 species (Chisholm et al., 2023; Davison, 2012), positioning it as a haven for concentrated biodiversity research. Comparatively, more studies can be emphasised on Singapore’s fungal biodiversity, with much of its ecology and functional properties remained unexplored, hampering efforts on further biotechnological utilisation, conservation, and understanding of the broader ecosystem (Choong, 2022; Lee & Choong, 2024; Weerakoon et al., 2015).

To address this imbalance, we implemented a rapid, streamlined and scalable solid-state fermentation workflow that bins collected isolates by terrestrial and marine origins and employed two general purpose media, identified for its ability to support robust biomass production across phylogenetically diverse lineages. Applied to a curated collection of 1,136 cryopreserved fungal isolates, this workflow achieved an 80.2% revival rate (n = 911; 99% CI: 77.1%-83.2%). High-resolution images were also collected across 3 key culture milestones [Day 7, Day 14, and pre-harvest (Day 11 to Day 71)] for some of the strains. Of the successful revived isolates, 94.1% (n = 857; 99% CI: 92.3%-95.9%) yielded sufficient biomass for DNA extraction, facilitating taxonomy curation with 18S. Quality assurances involved decontamination screening to omit non-fungal sequences and exclusion of fungal genomes lacking pre-harvest colony images.

The eventual curated collection comprises 518 diverse fungal strains, each paired with colony morphology images across various timepoints amounting to a total of 1,528 images. We processed the 518 fungal 18S rRNA sequences with multiple alignment using fast Fourier transform (MAFFT) to produce uniform base pair alignments (Katoh & Standley, 2013). With this alignment, we used FastTree to infer a maximum-likelihood phylogeny (Price et al., 2010), applied 100 bootstraps to obtain a majority-rule consensus tree, followed by visualisation with GGTREE (Yu et al., 2017). Annotated with taxonomic assignments and ecological metadata, the phylogenetic tree offers a comprehensive overview of Singapore’s fungal diversity. Spanning 127 genera and 3 unassigned strains across 16 classes and 4 phyla, the dataset represents fungal strains from a broad spectrum of habitats across the island, from soils and freshwater systems to marine environments (Figure 1; Table 1).

**Table 1.**
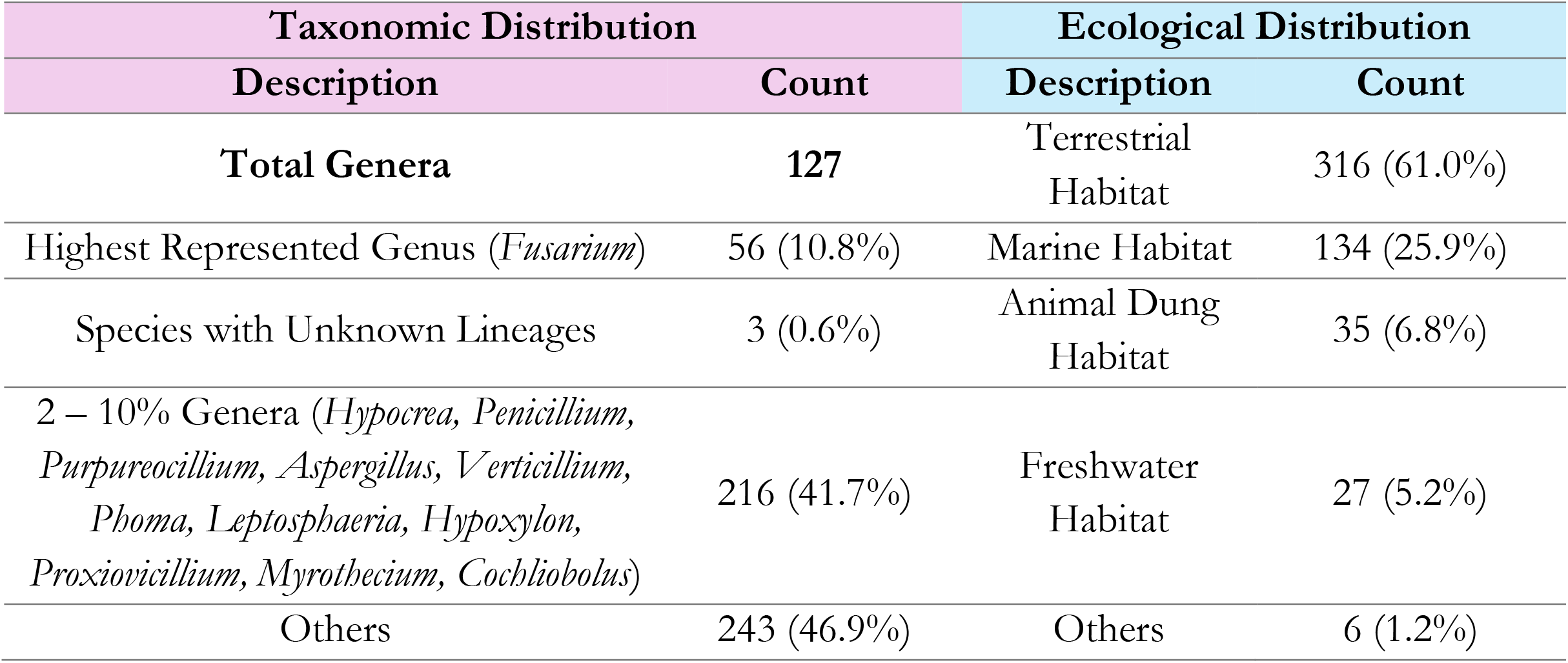
Taxonomic and Ecological Distribution of 518 Singapore Fungal Strains.

**Figure 1.**
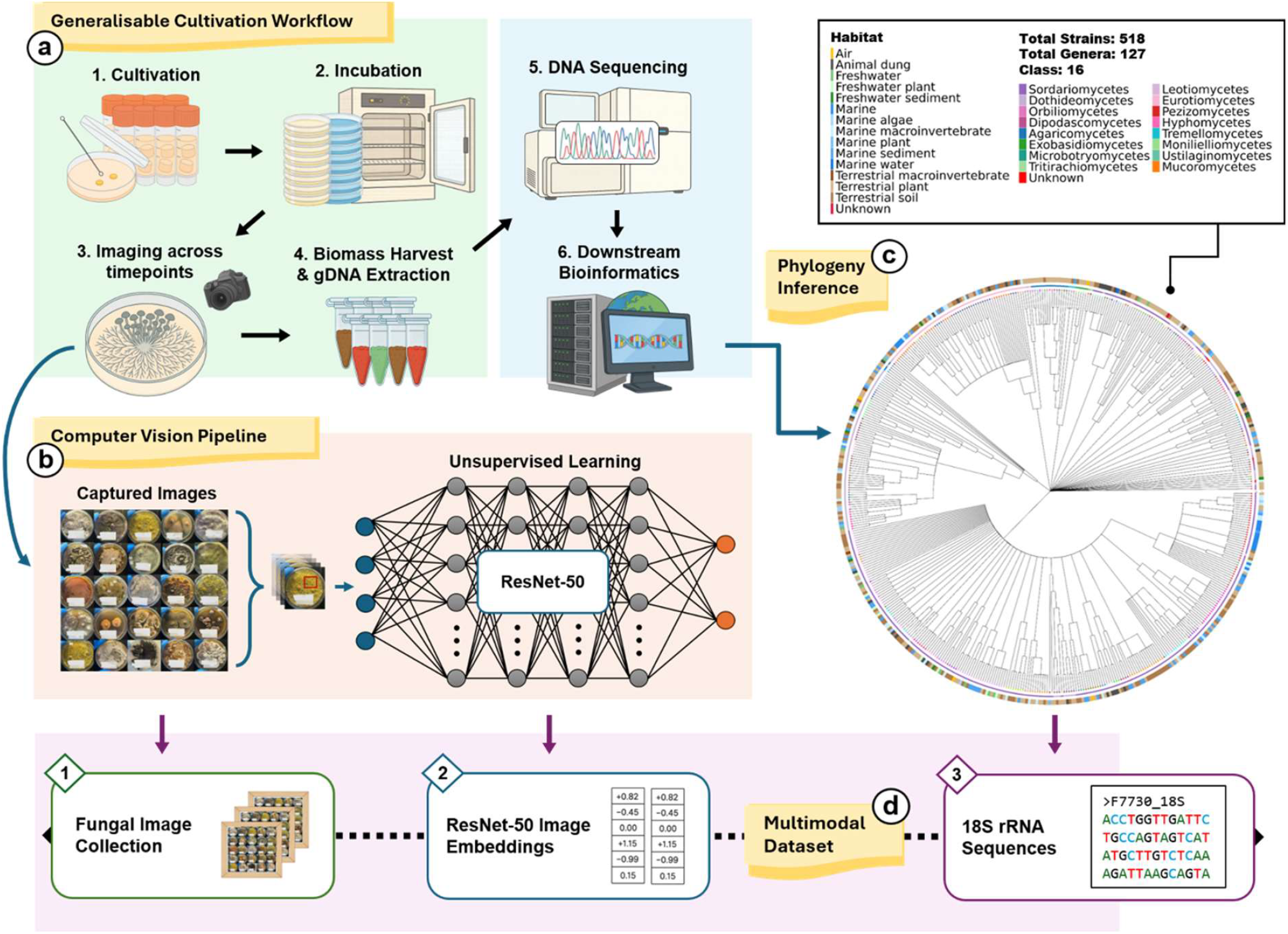
Overview of the multi-modal fungal dataset, providing an integrated perspective into the phenotypic and genomic diversity of Singapore’s fungi. (a) The cultivation workflow involves the following steps: First, (1) cryopreserved isolates are revived through solid-state cultivation and (2) incubated under standardised conditions, with (3) high-resolution colony images captured at key developmental milestones. Once colonies reach target biomass, samples underwent (4) DNA extraction, library preparation, and sequencing. (b) Computer vision pipeline for the processing of fungal colony images to extract high-dimensional feature embeddings. (c) Phylogenetic tree of 518 fungal strains annotated with class-level taxonomy and habitat origin. (d) Generated multimodal dataset consisting of (1) fungal image collection across key development milestones, (2) ResNet-50 embeddings of all images, and (3) 18S rRNA sequences for taxonomic inference.

To demonstrate the utility of the multi-modal dataset, we focused on 518 pre-harvest colony images drawn from the complete image collection of 1,528. High-dimensional representations of image features of the strain collection were extracted using a Residual Network with 50 layers (ResNet-50) convolutional neural network (CNN) (He et al., 2015). Prior to feature extraction, input images were pre-processed with the Efficient and Accurate Scene Text (EAST) detection model to identify and occlude text regions (Zhou et al., 2017), ensuring downstream analyses were driven solely by morphological traits. The resulting image embeddings effectively captured ecological niches, providing a direct linkage between phenotype and taxonomy. To evaluate the efficacies of the captured images and their corresponding embeddings in preserving taxonomic resolution and ecological niches, we projected the 2048-dimensional embeddings into two-dimensions using t-distributed stochastic neighbour embedding (t-SNE) (Figure 2). Overlaid ecological metadata and captured pre-harvest images, this projection (Figure 2a) revealed interesting phenotypic clusters, most notably by four coherent clusters defined by pigmentation: (2a-1) strains from diverse habitats exhibiting a deep black phenotype; (2a-2) clear white phenotypes from freshwater and marine sources; (2a-3) yellow strains predominantly from marine sources. Morphology-based clustering was also evident, highlighted by a cluster of strains displaying circular and lobate colony morphotype (Figure 2a-4). Alternative overlaying with taxonomical metadata enabled seamless toggling between phenotype and taxonomy context (Figure 2b). For example, the black phenotype cluster (Figure 2a-1) corresponds to Dothideomycetes genera such as *Hortaea* and *Cochliobolus* (Figure 2b-1), both documented for their dark colony morphology (Eliahu et al., 2007; Plemenitaš et al., 2008). This first-pass analysis validates our computer vision pipeline and further demonstrates the utility of the multimodal dataset to capture meaningful biological nuances.

**Figure 2.**
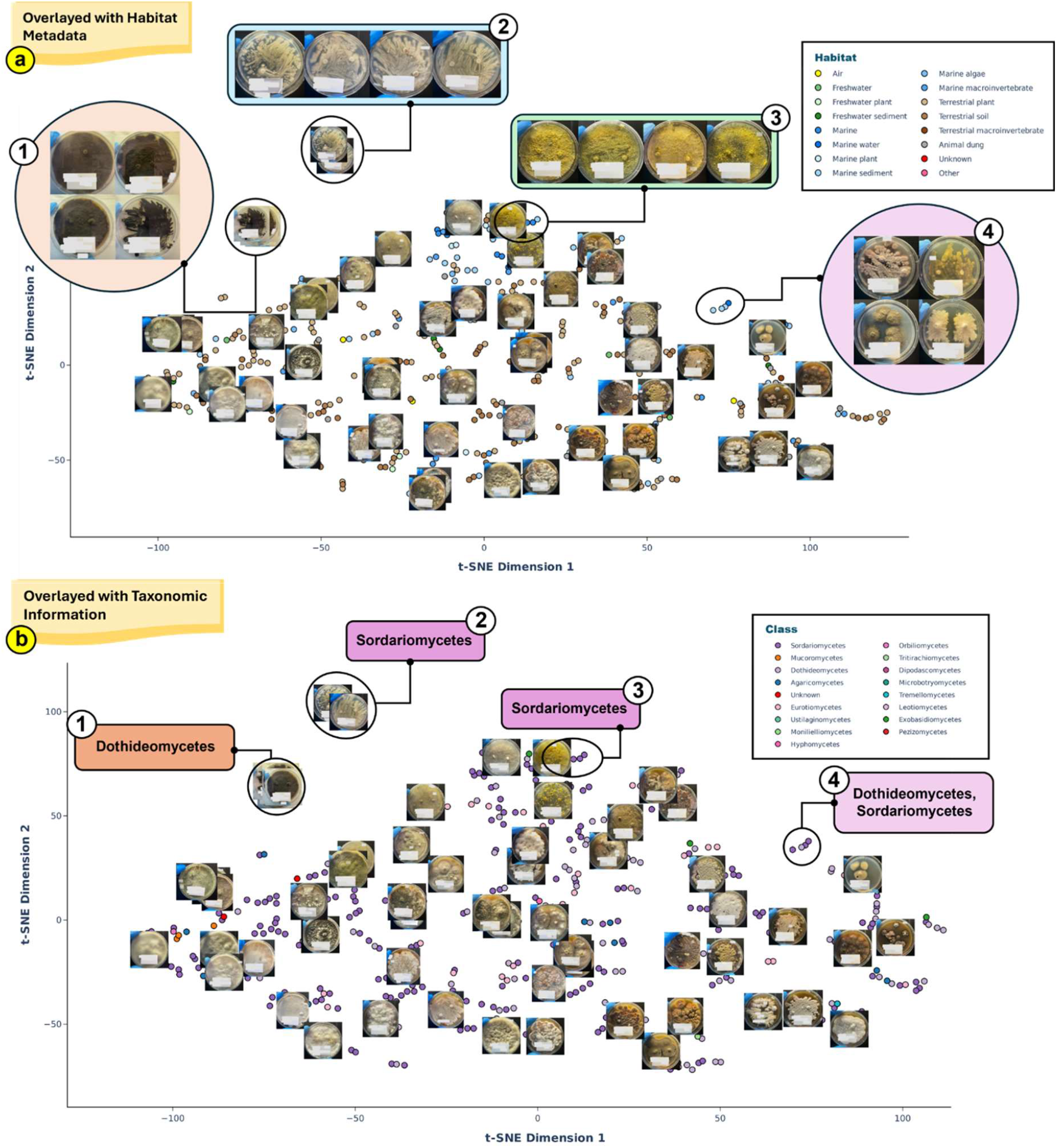
t-SNE projections of 518 ResNet-50 derived feature embeddings from pre-harvest fungal colony images, facilitating phenotypic-genomic analysis. Embeddings (points) are coloured with different metadata, facilitating phenotypic-genomic analysis. (a) Embeddings coloured according to the isolate’s habitat of origin, with emphasis on four key clusters with distinct morphotypes: (1) deep-black strains, (2) clear-white strains, (3) yellow strains, and (4) circular and lobate colony morphology. (b) The same embedding space as (a), coloured with class-level taxonomy, highlighting the same four clusters.

Beyond these exploratory findings, this dataset provides not only a streamlined cultivation workflow adaptable to a wide spectrum of fungal taxa but also presents a unique opportunity to interrogate the distinct colony morphology and underlying genomic signatures of 518 fungal strains from Singapore. It includes time-stamped fungal images across key developmental milestones, high-dimensional ResNet-50 embeddings, predicted 18S rRNA sequences, and corresponding taxonomy and ecological metadata. Together, these components form a unified platform for bridging visual phenotype to taxonomy, serving both as a reference for fungal biodiversity in tropical Singapore and as a hypothesis-generation platform, spurring targeted future investigations and supporting downstream genomic exploration for deep phenotypic and taxonomic insights.

## Methods

### Cultivation, Imaging and Harvest

A total of 1,136 previously uncharacterized fungal isolates, derived from Singapore, were cultivated. These strains are part of the Natural Product Library (NPL) collection in Agency for Science, Technology and Research (A*STAR) (Ng et al., 2018). Cryovials containing preservation fluid (reverse osmosis water with 25% glycerol content) and fungal agar plugs (agar with fungal mycelium) were retrieved from storage (−80°C freezer) and allowed to thaw once at room temperature. Cryopreserved fungal plugs were cultured on either ME [malt extract (30g/L), agar (15 g/L), mycological peptone (5 g/L)] or GYS [sea salt (40g/L), agar (15g/L), glucose (10g/L), yeast extract (1g/L)] agar, selected based on terrestrial versus marine origin. Cultures were incubated at 25 °C (42 °C for thermophilic isolates), and colony morphology was photographed in a biosafety cabinet across key developmental milestones (varied based on growth rate) using an iPhone 15 Pro (48-megapixel, macro mode).

### DNA Sequencing, Taxonomic Inference and Phylogenetic Analysis

Fungal DNA was extracted from biomass and sequenced on the Illumina NovaSeq 6000 platform. 18S regions were predicted with (Barrnap, v0.9), applying eukaryote-specific hidden Markov model (HMM) (Seemann, 2013). To facilitate genus-level taxonomic assignments, the predicted 18S regions of 518 strains were queried against NCBI core nucleotide database (accessed on 24 April 2025) with BLASTn (v2.14.1). Upon filtering by bit-score, each strain was provisionally assigned to the genus corresponding to its top BLAST hit, supplemented with phylum, sub-phylum, and class assignments with records from JGI’s MycoCosm (accessed on 4 September 2024) and MycoBank database (accessed on 1 June 2025).

To construct a phylogenetic tree, we performed multiple sequence alignment (MSA) on the 518 fungal 18S rRNA sequences using MAFFT (v7.525) executed with the --auto mode (FFT-NS-2 algorithm) (Katoh & Standley, 2013), The resulting alignment consisted of 518 sequences with equal length (2,287 bp). We generated 100 bootstrap replicates by resampling alignment columns with replacement and reconstructed maximum-likelihood trees with FastTree 2 (v2.1.11) (Price et al., 2010), employing the Generalized Time Reversible (GTR) model with gamma-distributed site-rate variation. Clade support values were derived by generating a majority-rule consensus tree using the R-based Analyses of Phylogenetics and Evolution package (ape, v5.8.1) (Paradis & Schliep, 2019), retaining clades supported by ≥50% of replicates (518 tips, 262 internal nodes, mean bootstrap support of 84%). The tree was subsequently visualised with GGTREE (Bioconductor v3.2) (Yu et al., 2017), enriched with genus, class, and habitat metadata.

### Image Pre-processing, ResNet-50, and t-SNE Visualisation

The 518 pre-harvest fungal colony images were first converted to .jpg format, then resized and padded (1024 pixel) while preserving original aspect ratio, to ensure dimensional compatibility with the frozen EAST text detection model (Zhou et al., 2017). The pretrained frozen EAST text detection model (inference-optimised) generated geometry maps and confidence scores for detected text presence, which were decoded into bounding boxes by translating model-predicted dimensions into input-image coordinates. Bounding boxes with low confidence score (<0.2) were discarded, retaining text regions within a generous confidence range. To mask these detected text regions and minimise bias in downstream phenotypic analysis, a Gaussian blur and semi-transparent overlay were applied to each bounding box. To facilitate inference and feature extraction by the ResNet-50 model, the masked images were first resized to a resolution of 224×224 pixels. They were then transformed into PyTorch tensors, converting pixel values to normalised floats in [Channel, Height, Width] format. These tensors were subsequently standardised using channel-wise mean [0.485, 0.456,

0.406] and standard deviation [0.229, 0.224, 0.225], aligning their input distribution with the ImageNet dataset, on which ResNet-50 was originally trained (He et al., 2015). Finally, the pre-processed image vectors were iteratively fed into the ResNet-50 model for unsupervised feature extraction, with the final average pooling layer of ResNet-50 CNN generating 2048-dimensional embeddings for each input images. Principal component analysis (PCA) and t-SNE, implemented via the scikit-learn (v1.6.1) suite, were applied to the 518 image embeddings, first denoising the original high-dimensional feature space to 10 principal components, followed by further dimensionality reduction to two-dimensions (n_components = 2, perplexity = 5, random state = 42) for effective visualisation. Relevant metadata (e.g., taxonomy, ecological origin) were overlaid onto the t-SNE plot to provide contextual interpretation of the ResNet-50-derived image clusters.

### Data Records

The 518 fungal dataset comprises of (1) fungal colony morphology images across key developmental stages (.jpg format), (2) ResNet-50 pre-harvest image embeddings (PyTorch format), (3) fungal strains taxonomy and ecological metadata, and (4) Barrnap-extracted 18S region (.fasta format). All components of the dataset have been deposited on Figshare and GitHub respectively.

### Technical Validation

Captured fungal colony images were first subjected to automated text occlusion by the frozen EAST text detection model, followed by processing with the ResNet-50 CNN model. The extracted 2048-dimensional embeddings capture ecological and taxonomical features, highlighting the robustness of the dataset.

### Usage Notes

This dataset enables users to investigate underexplored fungal lineages and help address gaps in the fungal phylogeny, with emphasis on Singapore’s rich mycobiota. Additionally, this dataset provides the unique opportunity to examine the dynamic colony morphology of 518 strains and explore its relationship to corresponding genomic signatures, presenting a valuable atlas for visual phenotype to taxonomic linkage. Accompanied by detailed strain metadata, the dataset, alongside Resnet-50 derived image embeddings and clustering, functions as a rapid hypothesis-generation platform. Users can identify fungal clusters of interest based on its phenotypic characteristics and rapidly relate its underlying genomics, creating a systematic and targeted exploration strategy for downstream functional or genomic studies. In addition, the phenotype–taxonomic framework can be expanded by integrating whole-genome context and is readily amenable to additional visual and language model applications, to extract deeper insights from the rich mine of visual and genomic data.

## Code Availability

Scripts for (1) 18S rRNA prediction with Barrnap, (2) BLASTn searches against the NCBI core nucleotide (core_nt) fungal database, (3) MAFFT alignment, (4) FastTree maximum-likelihood tree inference and (5) ResNet-50 embedding generation are available at https://github.com/twxdarren/fungal-trove.git.

## Acknowledgements

This work is supported by the Agency for Science, Technology and Research (A*STAR) under C233017006, and the Singapore Integrative Biosystems and Engineering Research Strategic Research & Translational Thrust (SIBER SRTT). The authors extend their gratitude towards Siew Bee Ng (A*STAR SIFBI) for her support on strains retrieval.

## Author Contributions

F.T.W., Y.H.L., and W.K. conceptualised, designed and coordinated the study. D.T.W.X conducted the experiments and data acquisition. D.T.W.X. wrote the manuscript along with inputs from all the authors.

## References

Ahrendt, S. R., Mondo, S. J., Haridas, S., & Grigoriev, I. V. (2023). MycoCosm, the JGI’s Fungal Genome Portal for Comparative Genomic and Multiomics Data Analyses. In F. Martin & S. Uroz (Eds.), Microbial Environmental Genomics (MEG) (Vol. 2605, pp. 271–291). Springer US. 10.1007/978-1-0716-2871-3_14

Aime, M. C., & Brearley, F. Q. (2012). Tropical fungal diversity: Closing the gap between species estimates and species discovery. Biodiversity and Conservation, 21(9), 2177–2180. 10.1007/s10531-012-0338-7

Bhattarai, K., Bhattarai, K., Kabir, M. E., Bastola, R., & Baral, B. (2021). Fungal natural products galaxy: Biochemistry and molecular genetics toward blockbuster drugs discovery. In Advances in Genetics (Vol. 107, pp. 193–284). Elsevier. 10.1016/bs.adgen.2020.11.006

Case, N. T., Berman, J., Blehert, D. S., Cramer, R. A., Cuomo, C., Currie, C. R., Ene, I. V., Fisher, M. C., Fritz-Laylin, L. K., Gerstein, A. C., Glass, N. L., Gow, N. A. R., Gurr, S. J., Hittinger, C. T., Hohl, T. M., Iliev, I. D., James, T. Y., Jin, H., Klein, B. S., … Cowen, L. E. (2022). The future of fungi: Threats and opportunities. G3 Genes|Genomes|Genetics, 12(11), jkac224. 10.1093/g3journal/jkac224

Chisholm, R. A., Kristensen, N. P., Rheindt, F. E., Chong, K. Y., Ascher, J. S., Lim, K. K. P., Ng, P. K. L., Yeo, D. C. J., Meier, R., Tan, H. H., Giam, X., Yeoh, Y. S., Seah, W. W., Berman, L. M., Tan, H. Z., Sadanandan, K. R., Theng, M., Jusoh, W. F. A., Jain, A., … Sin, Y. C. K. (2023). Two centuries of biodiversity discovery and loss in Singapore. Proceedings of the National Academy of Sciences, 120(51), e2309034120. 10.1073/pnas.2309034120

Choong, M. F. A. (2022). Education on plants and fungi in Singapore: An urgent call. Nature in Singapore, Supplement 1, 279286. 10.26107/NIS-2022-0128

Crous, P. W., Grams, W., Stalpers, J. A., Robert, V., & Stegehuis, G. (2004). MycoBank: An online initiative to launch mycology into the 21st century. Studies in Mycology, 50(1), 19–22.

Danner, C., Mach, R. L., & Mach-Aigner, A. R. (2023). The phenomenon of strain degeneration in biotechnologically relevant fungi. Applied Microbiology and Biotechnology, 107(15), 4745–4758. 10.1007/s00253-023-12615-z

Davison, G. (2012). Special Ecology Feature: Biodiversity’s Crucial Role in the Modern Singapore City. CITYGREEN, 01(04), 102. 10.3850/S2382581212010617

Eliahu, N., Igbaria, A., Rose, M. S., Horwitz, B. A., & Lev, S. (2007). Melanin Biosynthesis in the Maize Pathogen Cochliobolus heterostrophus Depends on Two Mitogen-Activated Protein Kinases, Chk1 and Mps1, and the Transcription Factor Cmr1. Eukaryotic Cell, 6(3), 421–429. 10.1128/EC.00264-06

Grigoriev, I. V., Nikitin, R., Haridas, S., Kuo, A., Ohm, R., Otillar, R., Riley, R., Salamov, A., Zhao, X., Korzeniewski, F., Smirnova, T., Nordberg, H., Dubchak, I., & Shabalov, I. (2014). MycoCosm portal: Gearing up for 1000 fungal genomes. Nucleic Acids Research, 42(D1), D699– D704. 10.1093/nar/gkt1183

He, K., Zhang, X., Ren, S., & Sun, J. (2015). Deep Residual Learning for Image Recognition (Version 1). arXiv. 10.48550/ARXIV.1512.03385

Huberman, L. B. (2021). Developing Functional Genomics Platforms for Fungi. mSystems, 6(4), 10.1128/msystems.00730-21. https://doi.org/10.1128/msystems.00730-21

Hyde, K. D. (2022). The numbers of fungi. Fungal Diversity, 114(1), 1–1. 10.1007/s13225-022-00507-y

Hyde, K. D., Baldrian, P., Chen, Y., Thilini Chethana, K. W., De Hoog, S., Doilom, M., De Farias, A. R. G., Gonçalves, M. F. M., Gonkhom, D., Gui, H., Hilário, S., Hu, Y., Jayawardena, R. S., Khyaju, S., Kirk, P. M., Kohout, P., Luangharn, T., Maharachchikumbura, S. S. N., Manawasinghe, I. S., … Walker, A. (2024). Current trends, limitations and future research in the fungi? Fungal Diversity, 125(1), 1–71. 10.1007/s13225-023-00532-5

Katoh, K., & Standley, D. M. (2013). MAFFT Multiple Sequence Alignment Software Version 7: Improvements in Performance and Usability. Molecular Biology and Evolution, 30(4), 772–780. 10.1093/molbev/mst010

Lee, S., & Choong, A. (2024). Checklist of Fungi Species with their Category of Threat Status for Singapore. In The Singapore red data book: Red lists of Singapore biodiversity (Third edition, pp. 515– 518). National Parks Board.

Ng, S. B., Kanagasundaram, Y., Fan, H., Arumugam, P., Eisenhaber, B., & Eisenhaber, F. (2018). The 160K Natural Organism Library, a unique resource for natural products research. Nature Biotechnology, 36(7), 570–573. 10.1038/nbt.4187

Paradis, E., & Schliep, K. (2019). ape 5.0: An environment for modern phylogenetics and evolutionary analyses in R. Bioinformatics, 35(3), 526–528. 10.1093/bioinformatics/bty633

Paterson, R. R. M., Solaiman, Z., & Santamaria, O. (2023). Guest edited collection: Fungal evolution and diversity. Scientific Reports, 13(1), 21438, s41598-023-48471–0. 10.1038/s41598-023-48471-0

Peay, K. G., Kennedy, P. G., & Talbot, J. M. (2016). Dimensions of biodiversity in the Earth mycobiome. Nature Reviews Microbiology, 14(7), 434–447. 10.1038/nrmicro.2016.59

Plemenitaš, A., Vaupotič, T., Lenassi, M., Kogej, T., & Gunde-Cimerman, N. (2008). Adaptation of extremely halotolerant black yeast Hortaea werneckii to increased osmolarity: A molecular perspective at a glance. Studies in Mycology, 61, 67–75. 10.3114/sim.2008.61.06

Price, M. N., Dehal, P. S., & Arkin, A. P. (2010). FastTree 2 – Approximately Maximum-Likelihood Trees for Large Alignments. PLoS ONE, 5(3), e9490. 10.1371/journal.pone.0009490

Pye, C. R., Bertin, M. J., Lokey, R. S., Gerwick, W. H., & Linington, R. G. (2017). Retrospective analysis of natural products provides insights for future discovery trends. Proceedings of the National Academy of Sciences, 114(22), 5601–5606. 10.1073/pnas.1614680114

Robert, V., Vu, D., Amor, A. B. H., Van De Wiele, N., Brouwer, C., Jabas, B., Szoke, S., Dridi, A., Triki, M., Daoud, S. B., Chouchen, O., Vaas, L., De Cock, A., Stalpers, J. A., Stalpers, D., Verkley, G. J. M., Groenewald, M., Dos Santos, F. B., Stegehuis, G., … Crous, P. W. (2013). MycoBank gearing up for new horizons. IMA Fungus, 4(2), 371–379. 10.5598/imafungus.2013.04.02.16

Seemann, T. (2013). barrnap 0.9: Rapid ribosomal RNA prediction. https://github.com/tseemann/barrnap

Stallman, J. K., Haelewaters, D., Koch Bach, R. A., Brann, M., Fatemi, S., Gomez-Zapata, P., Husbands, D. R., Jumbam, B., Kaishian, P. J., Moffitt, A., & Catherine Aime, M. (2024). The contribution of tropical long-term studies to mycology. IMA Fungus, 15(1), 35. 10.1186/s43008-024-00166-5

Tedersoo, L., Bahram, M., Põlme, S., Kõljalg, U., Yorou, N. S., Wijesundera, R., Ruiz, L. V., Vasco-Palacios, A. M., Thu, P. Q., Suija, A., Smith, M. E., Sharp, C., Saluveer, E., Saitta, A., Rosas, M., Riit, T., Ratkowsky, D., Pritsch, K., Põldmaa, K., … Abarenkov, K. (2014). Global diversity and geography of soil fungi. Science, 346(6213), 1256688. 10.1126/science.1256688

Weerakoon, G., Ngo, K. M., Lum, S., Lumbsch, H. T., & Lücking, R. (2015). On time or fashionably late for lichen discoveries in Singapore? Seven new species and nineteen new records of Graphidaceae from the Bukit Timah Nature Reserve, a highly urbanized tropical environment in South-East Asia. The Lichenologist, 47(3), 157–166. 10.1017/S0024282915000043

Wu, B., Hussain, M., Zhang, W., Stadler, M., Liu, X., & Xiang, M. (2019). Current insights into fungal species diversity and perspective on naming the environmental DNA sequences of fungi. Mycology, 10(3), 127–140. 10.1080/21501203.2019.1614106

Yu, G., Smith, D. K., Zhu, H., Guan, Y., & Lam, T. T. (2017). GGTREE: An R package for visualization and annotation of phylogenetic trees with their covariates and other associated data. Methods in Ecology and Evolution, 8(1), 28–36. 10.1111/2041-210X.12628

Zhou, X., Yao, C., Wen, H., Wang, Y., Zhou, S., He, W., & Liang, J. (2017). EAST: An Efficient and Accurate Scene Text Detector (Version 2). arXiv. 10.48550/ARXIV.1704.03155

